# Ciliary cAMP regulates Shh signal interpretation to drive polarisation of differentiating neurons

**DOI:** 10.64898/2026.03.11.711079

**Authors:** Gabriela Toro-Tapia, Holly B. Burbidge, Veronica Biga, John R. Davis, Raman M. Das

## Abstract

Cellular differentiation is characterised by transitions in cell-state through reinterpretation of extracellular signals. However, the mechanisms facilitating context-dependent signal interpretation remain poorly understood. Differentiating neurons remodel a molecularly distinct primary cilium as they switch from canonical to non-canonical Shh signalling. Here, using long-term live-tissue imaging, we demonstrate that the opposing Shh signalling modulators Smo and GPR161 simultaneously accumulate in the remodelled primary cilium. The correct balance of Smo and GPR161 leads to elevated ciliary cAMP levels, which suppresses Gli transcription factor activation and regulates actin dynamics to drive neuron polarisation. Notably, disrupting this balance through Smo hyperactivation or GPR161 depletion results in reduced ciliary cAMP, inappropriate activation of canonical Shh signalling, dysregulated actin dynamics and initiation of multiple unstable axon-like projections. Thus, this study identifies shifts in ciliary cAMP levels as a key regulator of cellular signal interpretation, and links primary cilium-mediated signal transduction to precise control of cytoskeletal organisation.

## Introduction

The developing vertebrate nervous system arises from neuroepithelial progenitor cells, which undergo cell divisions to generate mature neurons within the neural tube^1^. These progenitor cells possess a primary cilium that projects into the lumen of the neural tube^2^. This specialised organelle operates as a compartmentalised centre for signal transduction, physically separated from the cytoplasm while maintaining selective communication with it^3^. In neuroepithelial progenitor cells, the primary cilium facilitates transduction of canonical Gli transcription factor-dependent Shh signalling, which mediates dorsoventral patterning of the early neural tube and maintains these cells as cycling progenitors^4,5^. In this context, the Shh modulators Smo and GPR161 have been proposed to exert opposing effects in regulating canonical Shh signalling^6^. In epithelial cells cultured *in vitro*, basal ciliary accumulation of GPR161 in the absence of Shh ligands, and therefore absence of ciliary Smo, prevents activation of canonical Shh signalling by elevating ciliary cAMP levels. This promotes formation of Gli repressor forms to inhibit transcription of Shh target genes^6^. In the presence of Shh ligands, ciliary accumulation of activated Smo mediates removal of GPR161 from the primary cilium, enabling Gli transcription factor activation and canonical Shh signal transduction^7^. Thus, mutually exclusive ciliary accumulation of Smo or GPR161 regulates the activation of canonical Shh signalling; however, it remains unclear if a similar mechanism is also operating in the *in vivo* context of developing tissues.

In the developing spinal cord, as neuroepithelial progenitor cells begin neuronal differentiation, they delaminate from the neuroepithelium through the regulated process of apical abscission, which mediates dismantling of the primary cilium (**Figure 1A**)^8^. This step correlates with acute cessation of canonical, Gli transcription factor-mediated Shh signalling and leads to cell cycle exit^8^. Following this, delaminating newborn neurons remodel a new primary cilium and undergo a switch to Smo-dependent non-canonical Shh signal interpretation and a drop in Gli transcriptional activity^9^. The remodelled primary cilium is characterised by accumulation of GPR161 and Smo^9^, but it is not clear if this ciliary accumulation is mutually exclusive. Furthermore, the mechanistic basis through which primary cilium remodelling mediates suppression of Gli transcription factor activity in cells undergoing Smo-mediated non-canonical Shh signal transduction is unknown.

**Figure 1:**
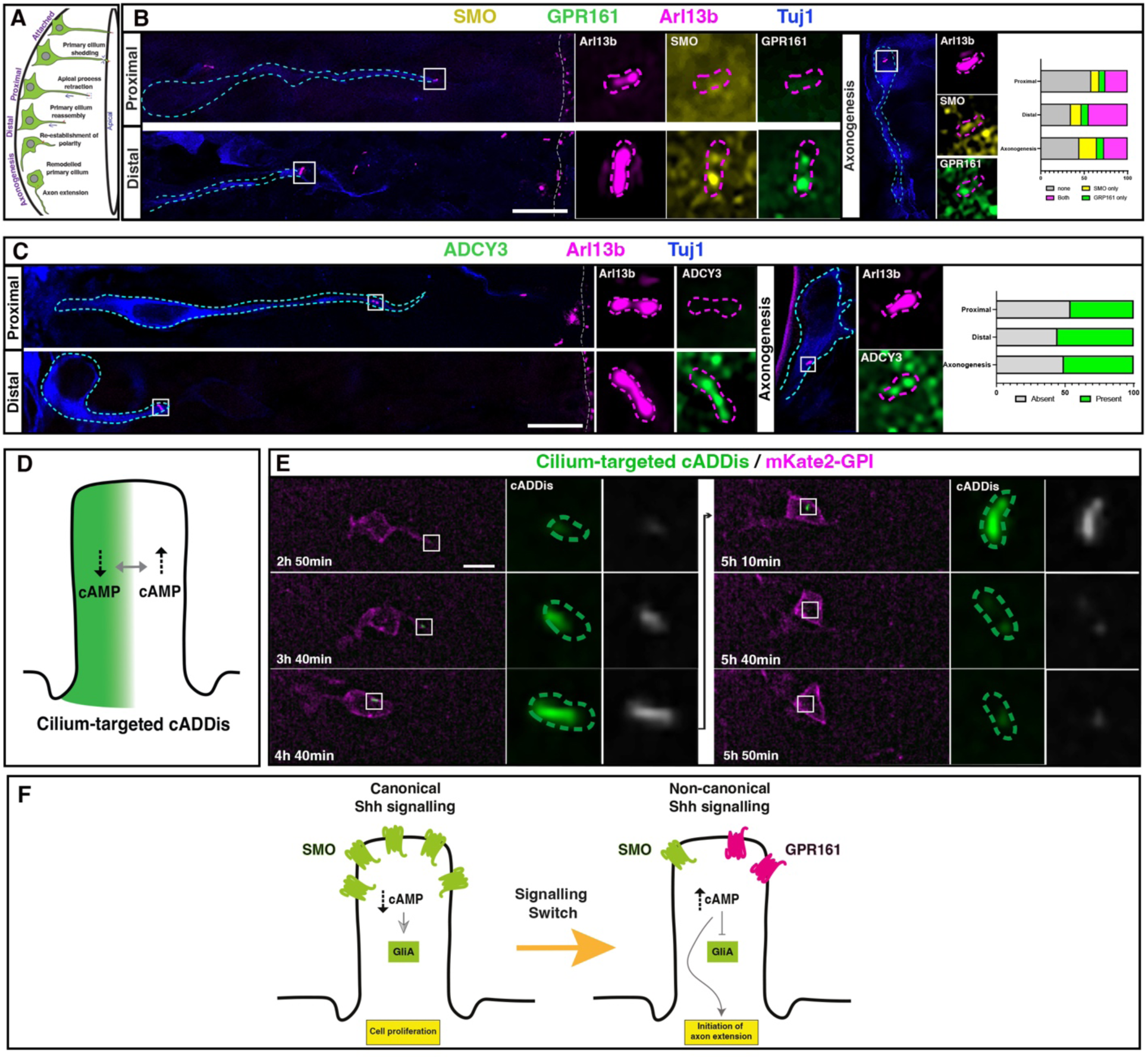
Simultaneous ciliary accumulation of Smo and GPR161 corresponds with elevated ciliary cAMP levels in differentiating neurons. **(A)** Representation of differentiating neurons at different stages of apical process retraction and early stages of axonogenesis in the developing spinal cord. **(B)** Immunostaining to detect endogenous Smo (yellow), GPR161 (green), Arl13b (magenta) and Tuj1(blue). Cyan dashed lines demarcate differentiating cells, magenta dashed lines demarcate primary cilia, and white boxes indicate enlarged regions to the right. Panel to the right displays quantification of Smo and GPR161 accumulation in primary cilia at different stages of apical process retraction and axon extension. **(C)** Immunostaining to label endogenous ADCY3 (green), Arl13b (magenta) and Tuj1 (blue). White boxes indicate enlarged regions to the right. Panel to the right displays quantification of ADCY3 accumulation in remodelled primary cilia at different stages of apical process retraction and axon extension. **(D)** Representation of the basis through which the cilium-targeted genetically encoded fluorescent cAMP sensor, cADDis, reports cAMP levels. cADDis fluorescence (green) decreases at high levels of ciliary cAMP. **(E)** Timelapse sequence of a differentiating neuron undergoing apical process retraction expressing cilium-targeted cADDis (green) and membrane marker mKate2-GPI (magenta). White boxes indicate enlarged regions to the right, and green dashed lines demarcate primary cilia. **(F)** Hypothetical mechanism of primary cilium mediated switch to non-canonical Shh signalling in differentiating neurons. Scale bars, 10μm.

Cells undergoing neuronal differentiation in the dorsal and intermediate neural tube undergo polarisation by initiating a single, ventrally directed axon^9^, a key step in the development of functional neural circuitry. Several signalling cues controlling outgrowth and navigation of axons that have initiated within the developing spinal cord have been identified^10^. Among these, Smo-mediated Gli transcription factor-independent Shh signalling has been shown to guide established axons towards the ventral midline of the developing neural tube^11^ by promoting cytoskeletal remodelling within the growth cone^12^. However, little is known about how context-specific Shh signal interpretation influences the cytoskeletal organisation of newborn neurons to direct initiation and maintenance of a single, stable axon.

Here, using long-term live tissue imaging, we reveal the mechanistic basis for the suppression of Gli transcription factor activity through primary cilium remodelling. We demonstrate that balanced simultaneous accumulation of the opposing Shh signalling modulators Smo and GPR161 elevates relative cAMP levels in the remodelled cilium to prevent Gli transcription factor activation, which corresponds with regulated actin dynamics that drive focused initiation of a single axon. This work positions ciliary cAMP as an orchestrator of cellular signal interpretation and links primary cilium-mediated signal interpretation with precise regulation of cytoskeletal organisation.

## Results

### Simultaneous ciliary accumulation of Smo and GPR161 corresponds with elevated ciliary cAMP levels in differentiating neurons

Remodelled primary cilia of differentiating neurons in the dorsal and intermediate chick neural tube display accumulation of the Shh modulators Smo and GPR161^9^. However, as it was not technically possible to perform co-labelling of Smo and GPR161 in chick tissue, it is not clear if these undergo concurrent ciliary accumulation. To investigate this, we utilised the improved fidelity of multi-label immunofluorescence in mouse embryos, which display a high degree of mechanistic conservation with chick during neuronal differentiation^13^. E10.5 mouse embryos were fixed and labelled for the early neuronal marker Tuj1, the primary cilium membrane marker Arl13b, Smo and GPR161 **(Figure 1B).** Following apical abscission, a minority of cells in the early stages of apical process retraction classified as proximal to the apical surface accumulated one (Smo only: 9%, 4/43 cells; GPR161 only: 7%, 3/43 cells, 4 embryos) or both of these Shh modulators (26%, 11/43 cells, 4 embryos, **Figure 1B, right panel**). In contrast, the majority of cells in the later stages of apical process retraction that were distal to the apical surface and had completed primary cilium reassembly exhibited ciliary accumulation of one or both of these Shh modulators (65%, 32/49 cells, 4 embryos), with 12% of cells accumulating Smo only, 8% accumulating GPR161 only, and a high proportion (45%) displaying accumulation of both Smo and GPR161. Most cells undergoing axon extension also accumulated these Shh modulators, but the proportion displaying simultaneous Smo and GPR161 accumulation was reduced (Smo and GPR161: 27%, 36/133 cells; Smo only: 20%, 27/133 cells; GPR161 only: 8%, 8/133 cells, 4 embryos). These observations reveal a high degree of simultaneous Smo and GPR161 accumulation in the remodelled primary cilia of cells in the later stages of apical process retraction, as they prepare to undergo axon initiation. Furthermore, the observed pattern of ciliary Smo and GPR161 accumulation is consistent with ciliary trafficking of these components and, consequently, further roles for these Shh modulators in regulating downstream effectors.

We hypothesised that the unexpected simultaneous Smo and GPR161 accumulation in the remodelled primary cilium of differentiating neurons may elevate ciliary cAMP levels and lead to repression of canonical Shh signalling. To investigate this, we first fixed and labelled E10.5 mouse embryos for the neuronal primary cilium-localised cAMP-producing enzyme adenylyl cyclase III (ADCY3)^14^ and Tuj1 (**Figure 1C**). This revealed ciliary accumulation of ADCY3 in approximately 50% of remodelled primary cilia at all stages of primary cilium reassembly (**Figure 1C, right panel**; proximal: 46%, n= 46/100 cells; distal: 55%, n= 46/81 cells; axonogenesis: 51% n=63/124 cells, 4 embryos). Therefore, GPR161 ciliary accumulation as neuronal differentiation progresses corresponds with accumulation of ADCY3, raising the possibility that ciliary GPR161 accumulation may facilitate elevated ciliary cAMP levels.

To investigate if ciliary GPR161 accumulation corresponds with elevated ciliary cAMP levels, we used a cilium-targeted version of the genetically encoded sensor for cAMP levels, 5HT6-cADDis (cilium-targeted cADDis)^15^. cADDis displays progressively diminished green fluorescence in response to increasing cAMP levels, facilitating direct visualisation of ciliary cAMP levels over time (**Figure 1D)**^16^. To confirm cADDis biosensor activity, chick embryo neural tubes, which are amenable for high-resolution live imaging^17^, were transfected with mRNA encoding for cilium-targeted cADDis. cADDis fluorescence levels were monitored in *ex-vivo* embryo slices cultured in medium containing 10 μM forskolin (Fsk), which directly activates adenylyl cyclases and increases basal cAMP levels **(Figure S1A-A’).** In contrast to cells imaged in medium containing carrier control DMSO, cells imaged in medium containing Fsk exhibited an abrupt decline in ciliary green fluorescence after 10 minutes of Fsk addition (Fsk, 16 cells; DMSO, 14 cells, 4 embryos), indicating a Fsk-mediated increase in ciliary cAMP levels **(Figure S1B-B’’).** We then monitored the dynamics of ciliary cAMP in cells undergoing neuronal differentiation by transfecting a DNA construct expressing mKate2-GPI to label cell membranes and mRNA encoding cilium-targeted cADDis (**Figure 1E, Movie S1**). This revealed high levels of green fluorescence in the reassembling primary cilia of newborn neurons undergoing apical process retraction, indicating low ciliary cAMP levels. However, as the remodelled primary cilia entered the neuronal cell body, a step that precedes axon initiation, we observed a sharp reduction of ciliary green fluorescence (10/10 cells, 10 embryos), indicating an increase in ciliary cAMP levels, which correlates with ciliary accumulation of GPR161. This observation introduced the possibility that elevated ciliary cAMP may facilitate suppression of canonical Shh signalling and drive initiation of axon extension (**Figure 1F)**.

### The correct balance of ciliary Smo and GPR161 accumulation facilitates elevated levels of ciliary cAMP in differentiating neurons

To investigate if GPR161 is required for initiation of axon extension, we performed RNA interference-mediated knockdown of GPR161 expression. Chick neuroepithelial cells were transfected with a mix of three short hairpin RNA-expressing constructs targeting different regions of GPR161 (pRFPRNAi-GPR161#1, pRFPRNAi-GPR161#2 and pRFPRNAi-GPR161#3), which also expressed RFP to facilitate labelling of transfected cells or a control firefly luciferase targeting construct (pRFPRNAi-Luc). Efficiency of knockdown was assessed by fixing embryos 48 hours after transfection and performing HCR RNA-FISH to detect GPR161 mRNA expression in transfected cells. Quantification of the number of HCR spots per cell revealed a significant reduction in GPR161 expression compared to cells transfected with pRFPRNAi Luc (**Figure S2;** GPR161 RNAi: 74 cells, 3 embryos; Luciferase RNAi: 65 cells, 3 embryos). The effect of GPR161 knockdown on neuron polarisation was then assessed by performing timelapse imaging of cells expressing the knockdown constructs and GFP-GPI to label cell membranes **(Figure 2**). We observed that while the majority of cells expressing pRFPRNAi Luc initiated a single axon and underwent normal axon extension towards the ventral midline (**Figure 2A, Movie S2**; 81% n=72/89 cells, 36 embryos), most cells depleted for GPR161 were unable to undergo normal axon initiation (**Figure 2A’, Movie S3**; 71%, n=78/110 cells, 38 embryos). A subset of these cells instead presented multiple short and transient axon-like projections, which did not mature into a stable axon and instead collapsed (29%, 32/110 cells). Further subsets either did not initiate axons (26%, n=28/110 cells) or extended axons in an incorrect orientation (16%, n = 18/110). Overall, these results indicate that GPR161 is required for correct neuron polarisation and axon initiation.

**Figure 2:**
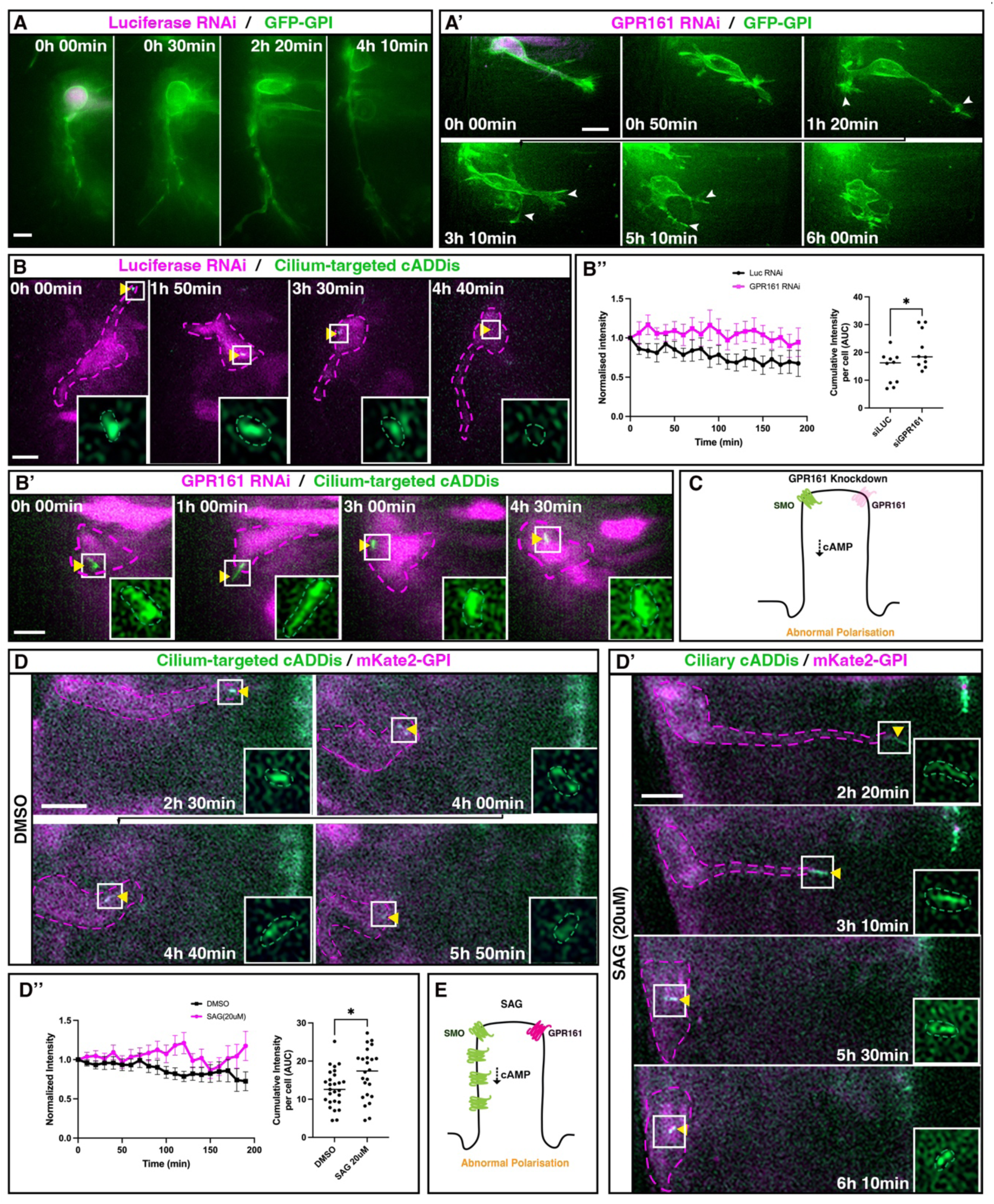
The correct balance of ciliary Smo and GPR161 accumulation facilitates elevated levels of ciliary cAMP in differentiating neurons. **(A)** Timelapse sequence of differentiating neuron expressing luciferase-targeting RNAi construct (magenta) and membrane marker GFP-GPI (green). **(A’)** Timelapse sequence of differentiating neuron expressing GPR161-targeting RNAi constructs (magenta) and GFP-GPI (green). White arrowheads indicate unstable axon-like projections. **(B)** Timelapse sequence of differentiating neuron expressing luciferase RNAi construct (magenta) and cilium-targeted cADDis (green). White boxes indicate enlarged regions in inset panels. Magenta dashed lines demarcate differentiating cells; green dashed lines demarcate primary cilia. **(B’)** Timelapse sequence of differentiating neuron expressing GPR161 RNAi constructs (magenta) and cilium-targeted cADDis (green). Yellow arrowhead indicates primary cilium. **(B’’)** Quantification of normalised relative cADDis fluorescence over 200 minutes in GPR161 knockdown (magenta line) and luciferase control cells (black line). Error bars represent standard error. Right-hand graph displays quantification of cumulative intensity per cell (AUC: area under the curve). Data analysed using unpaired *t*-test (**p**=0.0297). **(C)** Schematic summarising the effect of GPR161 knockdown on relative ciliary cAMP levels. **(D)** Timelapse sequence of differentiating neuron expressing cilium-targeted cADDis (green) and mKate2-GPI (magenta) imaged in medium containing DMSO. **(D’)** Timelapse sequence of differentiating neuron expressing cilium-targeted cADDis (green) and mKate2-GPI (magenta) imaged in medium containing 20 μM SAG. **(D’’)** Quantification of normalised relative cADDis fluorescence over 200 minutes in SAG-treated (magenta line) and DMSO-treated (black line) cells. Error bars represent standard error. Right-hand graph displays quantification of cumulative intensity per cell (AUC, area under the curve). Data analysed using an unpaired *t*-test (**p**=0.0213). **(E)** Schematic summarising the effect of SAG on relative ciliary cAMP levels. Scale bars: 10μm.

The effect of GPR161 knockdown on ciliary cAMP levels was then monitored by transfecting cells with the knockdown constructs and cilium-targeted cADDis followed by timelapse imaging for 5 hours. Consistent with our earlier observation (**Figure 1E),** control cells expressing pRFPRNAi-Luc displayed a decline in relative levels of cADDis fluorescence as the remodelled primary cilium entered the neuronal cell body, indicating elevated cAMP levels (**Figure 2B and B’’, Movie S4**; 10/10 cells, 9 embryos). Conversely, in cells expressing the GPR161 knockdown constructs, relative levels of cilium-localised cADDis fluorescence were maintained as the remodelled primary cilium entered the cell body, indicating low levels of ciliary cAMP (**Figure 2B’ and B’’, Movie S5**; 10/10 cells, 8 embryos). These results suggest that GPR161 accumulation in the remodelled primary cilium leads to elevated ciliary cAMP levels in differentiating neurons undergoing polarisation (**Figure 2C**).

Smo and GPR161 are suggested to have opposite effects on cAMP production. While Smo ciliary accumulation has been proposed to decrease ciliary cAMP, allowing activation of Gli transcription factors and canonical Shh signalling^15^, ciliary accumulation of GPR161 is proposed to elevate ciliary cAMP, which negatively regulates canonical Shh signalling by promoting processing of Gli transcriptional factors to their repressor forms^6,18^. Given that the remodelled primary cilium accumulates both Smo and GPR161, knockdown of GPR161 likely creates an imbalance between these two modulators of the Shh signalling response, leading to increased relative levels of Smo and reduced ciliary cAMP levels. We therefore hypothesised that the balance between ciliary Smo and GPR161 may determine appropriate cAMP levels in the remodelled primary cilium.

To test this hypothesis, we hyperactivated Smo using the Smoothened agonist SAG, which promotes activation and enhances ciliary accumulation of Smo^19^, thus tipping the balance in favour of Smo (**Figure 2E)**. Cells were transfected with cilium-targeted cADDis and mKate2-GPI and then subjected to timelapse imaging in medium containing 20μM SAG or DMSO carrier control. Cells imaged in medium containing DMSO typically displayed a sharp increase in ciliary cAMP levels as the remodelled primary cilium entered the cell body (**Figure 2D and D’’, Movie S6**; 27 cells, 13 embryos). In contrast, cells imaged in medium containing SAG displayed low levels of ciliary cAMP over time (**Figure 2D’ and D’’, Movie S7**; 25 cells, 13 embryos). In addition, these cells were unable to initiate axon extension, similar to GPR161 knockdown cells **(Figure 2A’)**. Overall, these results strongly suggest that the correct balance of ciliary Smo and GPR161 regulates ciliary cAMP levels, leading to normal neuron polarisation and axon initiation.

### Ciliary cAMP levels modulate Gli transcription factor activation

To determine if elevated ciliary cAMP in polarising neurons facilitates the switch to non-canonical Shh signalling through suppression of Gli activation, we used a reporter of Gli-mediated transcription GBS-NLS-KikGR, which carries eight copies of the Gli binding site from the FoxA2 enhancer^20^ upstream of the green-to-red photoconvertible fluorescent protein KiKGR fused to a nuclear localisation signal (NLS). Following photoconversion of the existing pool of KiKGR, this reporter facilitated monitoring of de-novo Gli transcriptional activity by monitoring green fluorescence levels (**Figure 3A**). To verify reporter activity, cells were transfected with mKate2-GPI and GBS-NLS-KikGR and subjected to live imaging. Consistent with previous work^9^, following photoconversion, we observed recovery of green nuclear KikGR expression in the progeny of mitotic cells (**Figure S3A, Movie S8;** 18/18 cells, 9 embryos), but not in differentiating neurons undergoing axon extension, which have undergone the switch to Gli-independent Shh signalling **(Figure S3B, Movie S9**; 12/12 cells, 11 embryos). We then used this reporter to monitor the mode of Shh signalling in cells that had undergone RNAi-mediated depletion of GPR161. Cells expressing control pRFPRNAi-Luc correctly initiated axon extension and, following photoconversion, did not recover green KikGR expression, indicating that these cells had undergone cessation of Gli-transcription dependent Shh signalling (**Figure 3B, Movie S10;** 12/14 cells, 10 embryos). Conversely, cells expressing pRFPRNAi-GPR161 were unable to mature an axon and displayed rapid recovery of green KikGR expression within 180 minutes on average (SD=139.9) (**Figure 3B’, Movie S11**; 14/16 cells,12 embryos). These results demonstrate that depletion of GPR161 results in inappropriate activation of Gli transcription factors and maintenance of canonical Shh signalling. These findings therefore suggest that relatively reduced ciliary cAMP levels, caused by shifting the ciliary Smo:GPR161 ratio towards Smo, promote activation of Gli transcription factors and maintenance of canonical Shh signalling (**Figure 3C**). To validate this, we increased ciliary Smo accumulation by applying SAG to cells expressing mKate2-GPI and GBS-NLS-KikGR (**Figure 3D**). Here, control cells imaged in medium containing DMSO extended axons normally and did not recover green KikGR expression following photoconversion (**Figure 3D**, **Movie S12**; 16/17 cells undergoing normal axon extension, 12 embryos). In contrast, similar to cells depleted for GPR161, cells imaged in medium containing SAG were unable to mature an axon and displayed recovery of green KikGR following photoconversion within 200 mins on average (SD=135.7) (**Figure 3D’ and E, Movie S13,** 15/21 cells, 20 embryos). Overall, these experiments suggest that the correct balance of ciliary Smo and GPR161 accumulation regulates relative ciliary cAMP levels to facilitate accurate interpretation of Shh signalling in differentiating neurons.

**Figure 3:**
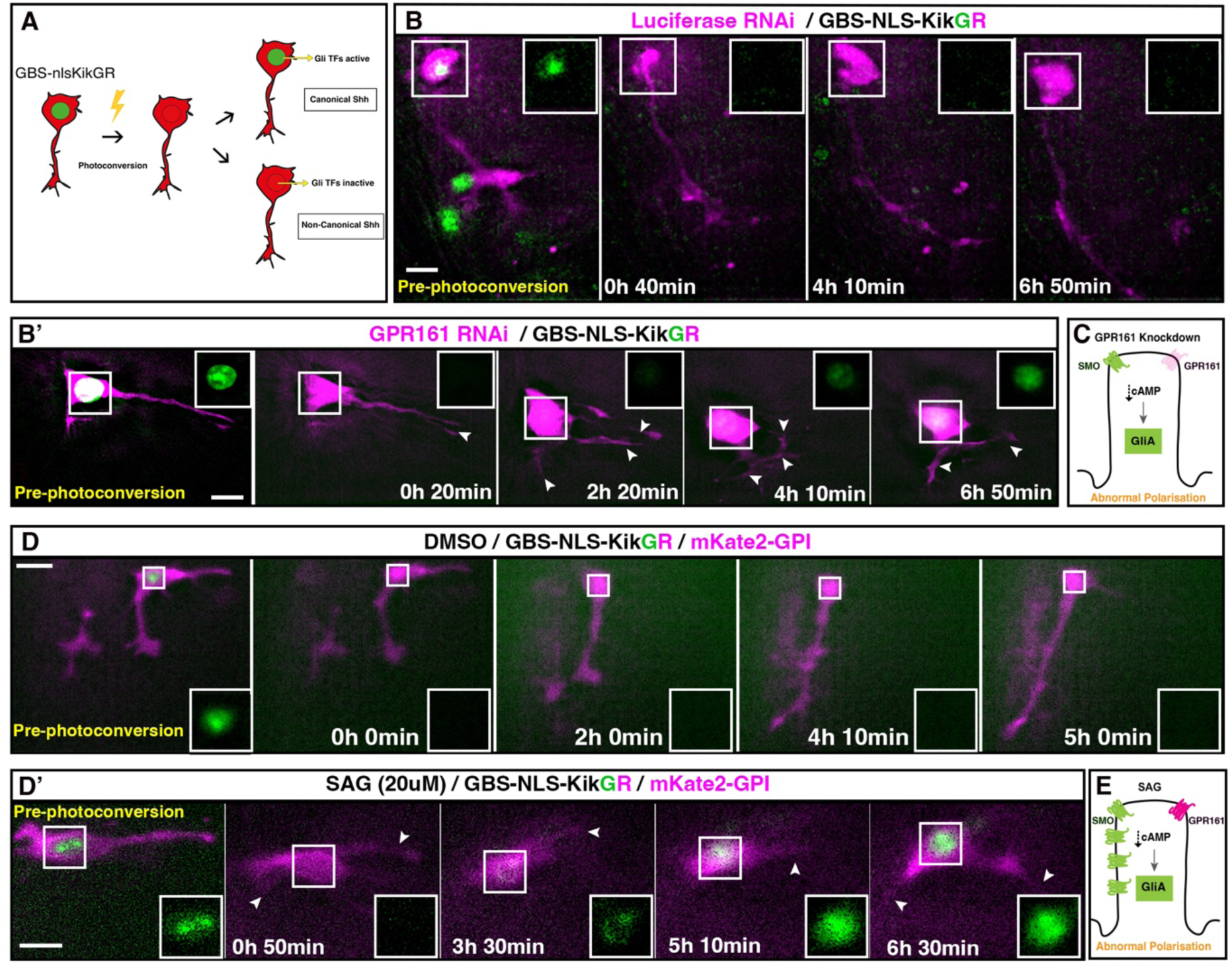
Ciliary cAMP levels modulate Gli transcription factor activation. **(A)** Green-to-red photoconvertible reporter for Gli transcriptional activity GBS-NLS-KikGR. Following photoconversion, nuclear GFP expression indicates activation of canonical Shh signalling. **(B)** Timelapse sequence of differentiating neuron expressing luciferase RNAi construct (magenta) and GBS-NLS-KikGR. First panel shows the cell before photoconversion. White boxes indicate regions enlarged in inset panels. **(B’)** Timelapse sequence of differentiating neuron expressing GPR161 RNAi constructs (magenta) and GBS-NLS-KikGR. **(C)** Schematic summarising the effect of GPR161 knockdown on relative ciliary cAMP levels and the mode of Shh signalling. **(D)** Timelapse sequence of differentiating neuron expressing GBS-NLS-KikGR and membrane marker mKate2-GPI (magenta) imaged in medium containing DMSO. **(D’)** Timelapse sequence of differentiating neuron expressing GBS-NLS-KikGR and mKate2-GPI (magenta) imaged in medium containing 20 μM SAG. **(E)** Schematic summarising the effect of SAG on relative ciliary cAMP levels and the mode of Shh signalling. Scale bars: 10μm.

### Primary cilium-mediated Shh signal interpretation facilitates normal actin dynamics in differentiating neurons

Neuron polarisation and axon extension involve highly dynamic remodelling of filamentous actin (F-actin), as accumulation of F-actin determines the site of axon initiation and stabilises axon extension^21–26^. Given our previous finding that disruption of primary cilium-mediated Shh signalling impedes neuron polarisation and axon extension^9^, we examined whether this was through dysregulation of normal actin dynamics. To investigate this, we first determined the normal actin dynamics that characterise axon initiation in the developing spinal cord. Cells were transfected with GFP-GPI and F-Tractin-mKate2 to label F-actin and subjected to timelapse imaging. During polarisation of differentiating neurons, we observed distinct regions of membrane-associated F-actin accumulations within the cell bodies of polarising neurons. These actin accumulations were localised in an apparently random fashion around the circumference of the cell body and were the sites of occasional transient filopodia and small, transient protrusions which did not develop into an extending axon. Consistent with previous reports^22,27^, we observed that one of these accumulations, mainly located towards the baso-ventral region of the cell body, subsequently matured into the emerging growth cone, which then continued to extend towards the ventral region of the spinal cord (**Figure 4A**, F-Tractin channel highlighted in **Figure S4A, Movie S14**; 22/25 cells, 17 embryos). These observations indicated that axon initiation is preceded by a clear accumulation of F-actin at the site of the emerging growth cone. Furthermore, in cells that had initiated axons, we observed that axon extension was characterised by dynamic actin accumulations along the axonal shaft **(Figure 4B**, F-Tractin channel highlighted in **Figure S4B, Movie S15**; 41/45 cells, 24 embryos). Similar to the actin accumulations observed in the cell bodies of polarising neurons, these axonal actin accumulations were dispersed along the axonal shaft and were often the site for transient filopodia or small, transient protrusions. To investigate these further, we performed super-resolution structured illumination timelapse imaging, which confirmed that axonal actin accumulations extended into axonal filopodia or small protrusions **(Figure S5**, **Movie S16,** 3/3 cells, 1 embryo), suggesting that actin polymerisation at these sites drives formation of transient cellular protrusions.

**Figure 4:**
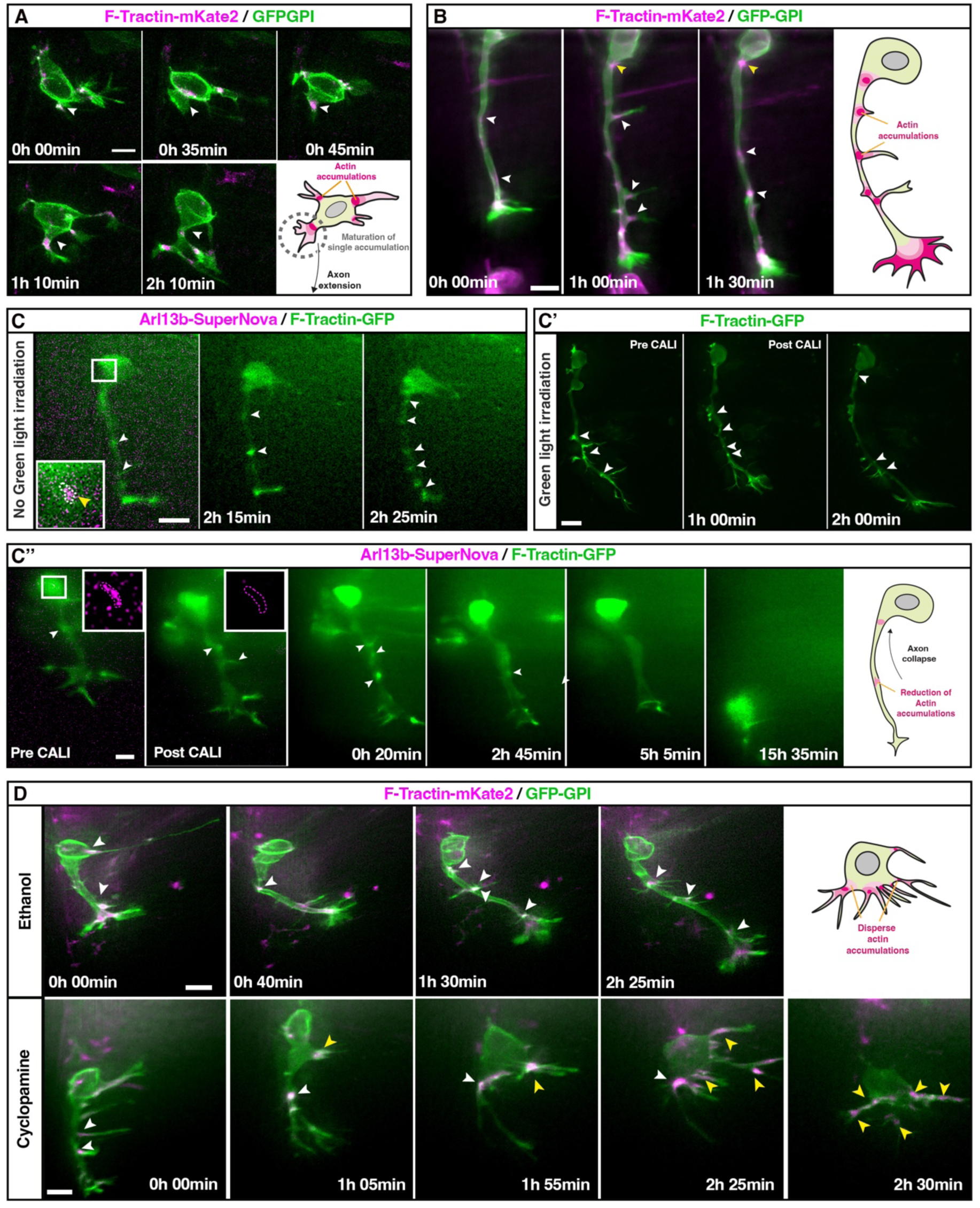
Primary cilium-mediated Shh signal interpretation facilitates normal actin dynamics in differentiating neurons. **(A)** Timelapse sequence of differentiating neuron expressing F-actin probe F-Tractin-mKate2 (magenta) and membrane marker GFP-GPI (green) undergoing axon initiation. White arrowheads indicate maturation of single actin accumulation into initiating axon. Bottom right schematic summarises actin dynamics in cells undergoing axon initiation. **(B)** Timelapse sequence of differentiating neuron expressing F-Tractin-mKate2 (magenta), and GFP-GPI (green) undergoing axon extension. White arrowheads indicate dynamic actin accumulations along the axon shaft. Right-hand schematic summarises actin dynamics in a cells undergoing axon extension. **(C)** Timelapse sequence of differentiating neuron expressing Arl13b-SuperNova (magenta) and F-Tractin-GFP (green) not subjected to green-light irradiation undergoing normal axon extension. White box indicates region enlarged in inset panel. White arrowheads indicate characteristic actin accumulations along the axon shaft. **(C’)** Timelapse sequence of differentiating neuron expressing F-Tractin-GFP (green) only undergoing normal axon extension following green light irradiation. **(C’’)** Timelapse sequence of differentiating neuron expressing Arl13b-SuperNova (magenta) and F-Tractin-GFP (green) following CALI-mediated disruption of primary cilium. First panel shows cell before green light irradiation (pre-CALI). Right-hand schematic summarises actin dynamics in differentiating neurons undergoing axon extension following CALI-mediated primary cilium disruption. **(D)** Timelapse sequences of differentiating neurons expressing F-Tractin-mKate2 (magenta), and GFP-GPI (green) undergoing axon extension imaged in medium containing ethanol (top panels) or cyclopamine (bottom panels). Top-right schematic summarises actin dynamics in cells imaged in medium containing 15 μM cyclopamine. White arrowheads indicate retained actin accumulations; yellow arrowheads indicate dispersed actin accumulations throughout the cell body. Scale bars: 10μm.

To establish a role for primary cilium-mediated signalling in directing these actin dynamics, we disrupted ciliary function using chromophore-assisted light inactivation (CALI) by fusing Arl13b to the phototoxic red fluorescent protein SuperNova, which locally releases reactive oxygen species in a confined radius lower than the average protein-protein interaction distance ^28^. We have previously demonstrated that this technique results in depletion of ciliary Arl13b and leads to collapse of extending axons^9^. Control cells transfected with Arl13b-SuperNova and F-Tractin-GFP but not subjected to green-light irradiation (**Figure 4C**, F-Tractin channel highlighted in **Figure S4C, Movie S17**; 9/10 cells, 7 embryos) or those transfected with F-Tractin-GFP only and subjected to green-light irradiation (**Figure 4C’,** F-Tractin channel highlighted in **Figure S4C’, Movie S18**; 8/8 cells, 7 embryos) exhibited normal actin dynamics along the axon shaft and correct axon extension. Green light irradiation of cells transfected with Arl13b-SuperNova and F-Tractin-GFP that were undergoing axon extension resulted in loss of cilium-localised red fluorescence, indicating disruption of ciliary Arl13b. Following this, we observed a gradual loss of axonal actin accumulations over 5-7 hours, which was subsequently accompanied by a marked collapse of the extending axon (**Figure 4C’’,** F-Tractin channel highlighted in **Figure S4C’’, Movie S19**; 9/12 cells, 11 embryos). These results suggest that signal transduction through the remodelled primary cilium facilitates the characteristic actin dynamics observed in cells undergoing axon extension.

We have previously demonstrated that disruption of Shh signalling using the Smo antagonist cyclopamine impairs axon extension^9^. To examine if this was through disrupted actin dynamics, we imaged cells expressing F-Tractin-mKate2 and GFP-GPI in medium containing cyclopamine. Cells imaged in medium containing carrier control ethanol underwent normal axon extension (**Figure 4D**, F-Tractin channel highlighted in **Figure S4D, Movie S20**; 21/25 cells, 12 embryos) with clear axonal actin accumulations associated with transient axonal protrusions. In contrast, most cells imaged in medium containing cyclopamine were unable to initiate an axon or collapsed axons that had already initiated (**Figure 4D**, F-Tractin channel highlighted in **Figure S4D, Movie S21;** 18/25 cells, 10 embryos). Here, cells that were unable to initiate axons or which underwent axon collapse displayed dispersed actin accumulations throughout the cell body (12/18 cells, 13 embryos), which were associated with multiple long, thin and dynamic filopodia-like protrusions (10/18 cells, 13 embryos). Therefore, inhibition of Smo function leads to small, dispersed actin accumulations around the cell body and formation of excessive thin transient protrusions, and impaired polarisation and axon initiation in differentiating neurons (**Figure 4D, top right panel**). Overall, these observations suggest that primary cilium-mediated Shh signal interpretation regulates actin dynamics to establish a stable axon.

### Ciliary cAMP levels direct actin dynamics to drive neuronal differentiation

We next investigated the possibility that suppression of canonical Shh signalling through elevated ciliary cAMP levels influences actin dynamics to drive initiation of a single stable axon. We first disrupted the balance between Smo and GPR161 by knocking down GPR161 and monitored F-actin dynamics by labelling with F-Tractin-GFP. Control cells expressing pRFPRNAi-Luc initiated a single stable axon and displayed normal axonal actin dynamics (**Figure 5A**, F-Tractin channel highlighted in **Figure S6A Movie S22**; 13/18 cells, 14 embryos). In contrast, cells depleted for GPR161 initiated multiple unstable axon-like projections that underwent rapid collapse as described in previous sections. Notably, these unstable axon-like projections emerged from aggregated actin accumulations within the cell body, leading to maturation of multiple actin clusters into transient protrusions. Additionally, these axon-like projections did not exhibit the dynamic axonal actin accumulations characteristic of normal axon extension. (**Figure 5A’,** F-Tractin channel highlighted in **Figure S6A’, Movie S23;** 18/19 cells, 11 embryos). These results strongly suggest that suppression of canonical Shh signalling through elevated ciliary cAMP levels modulates the dynamics of F-actin to facilitate stable initiation of a single axon. To confirm this, we reduced ciliary cAMP levels by application of the Smo agonist SAG. Cells imaged in medium containing DMSO carrier control initiated a single axon from a distinct actin accumulation located in the baso-ventral region of the cell body, which then continued to extend and displayed normal axonal actin dynamics (**Figure 5B**, F-Tractin channel highlighted in **Figure S6B, Movie S24,** 34/49**, 17 embryos**). Similar to GPR161 knockdown cells, cells imaged in medium containing SAG initiated multiple unstable axon-like projections from regions of the cell body that displayed aggregated actin accumulations (**Figure 5B’,** F-Tractin channel highlighted in **Figure S6B’, Movie S25**; cells, 25/30, 15 embryos). These transient axon-like projections also did not display the normal dynamic actin accumulations that characterise stably extending axons.

**Figure 5:**
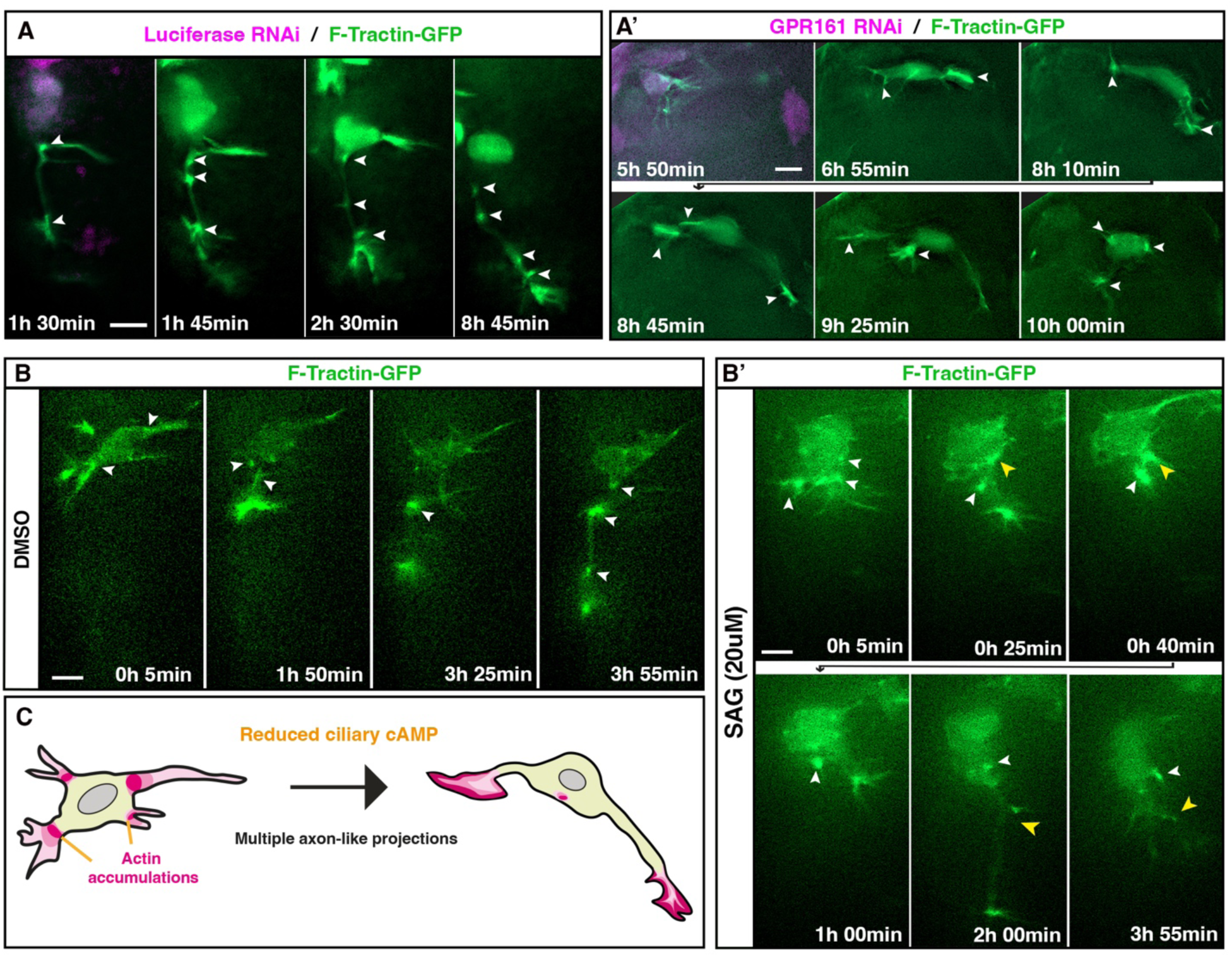
Ciliary cAMP levels direct actin dynamics to drive neuronal differentiation. **(A)** Timelapse sequence of differentiating neuron expressing luciferase RNAi construct (magenta) and F-Tractin-GFP (green) to label filamentous actin. White arrowheads indicate dynamic actin accumulations in axon shaft. **(A’)** Timelapse sequence of differentiating neuron expressing GPR161 RNAi constructs (magenta) and F-Tractin-GFP (green). White arrowheads indicate actin aggregation in unstable axon-like projections. **(B)** Timelapse sequence of differentiating neuron expressing F-Tractin-GFP (green) imaged in medium containing DMSO. White arrowheads indicate dynamic axonal actin accumulations. **(B’)** Timelapse sequence of differentiating neuron expressing F-Tractin-GFP (green) imaged in medium containing 20 μM SAG. White arrowheads indicate actin aggregations in cell body. Yellow arrowheads indicate initiating unstable axon-like projections. **(C)** Schematic summarising the effect of reduced ciliary cAMP on actin dynamics and maturation of multiple axon-like projections. Scale bars: 10μm.

Therefore, manipulations that reduce relative ciliary cAMP levels lead to the maturation of multiple aggregated F-actin accumulations, which subsequently develop into growth cone-like structures and give rise to multiple transient axon-like projections. Taken together, these findings indicate that elevated ciliary cAMP levels and suppression of canonical Shh signalling is crucial for regulation of the dynamics and spatial organisation of actin during neuronal differentiation. This leads to the controlled initiation and maturation of a single, distinct axon, which emerges from a baso-ventral actin accumulation in the cell body and maintains multiple dynamic F-actin accumulations along the axon shaft (**Figure 6**).

**Figure 6:**
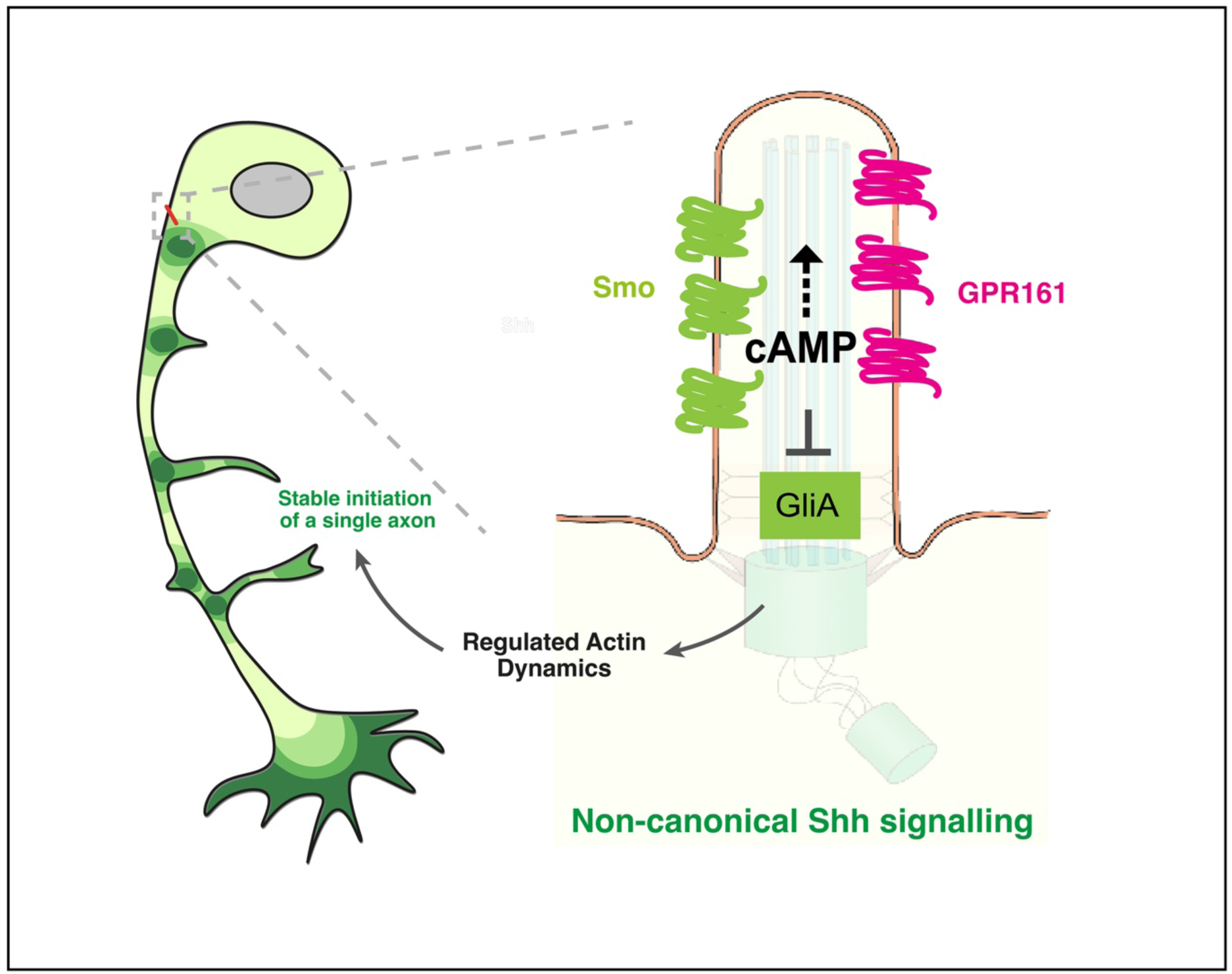
Ciliary cAMP levels mediate a switch in Shh signalling to drive polarisation of differentiating neurons. The correct balance of GPR161 and Smo ciliary accumulation modulates the switch to non-canonical Shh signalling through elevated ciliary cAMP levels to regulate the dynamics and spatial organisation of actin in differentiating neurons. The correct levels of ciliary cAMP leads to initiation and maturation of a single stable axon, which emerges from a baso-ventral actin accumulation in the cell body and maintains multiple dynamic F-actin accumulations along the extending axonal shaft.

## Discussion

This work emphasises the role of the primary cilium in facilitating accurate, context-specific interpretation of extracellular signals *in vivo* and demonstrates how regulation of signal interpretation shapes the actin cytoskeleton to drive initiation of a single axon during neuronal polarisation *in vivo*. We demonstrate that the remodelled primary cilium displays a unique molecular configuration, characterised by simultaneous accumulation of the Shh signalling modulators Smo and GPR161. This is notable, as in the context of canonical Shh signal transduction, ciliary accumulation of GPR161 suppresses Gli activation, thereby repressing Smo-mediated Shh signalling^6,18^. However, differentiating neurons require active, Smo-mediated non-canonical Shh signalling to undergo correct polarisation and axon initiation^9^. Here, we reveal that the correct balance of ciliary Smo and GPR161 in polarising neurons leads to elevated ciliary cAMP levels, which prevent Gli transcription factor activation to drive the switch to Gli-independent Shh signalling. Furthermore, our findings suggest that elevated ciliary cAMP levels regulate actin dynamics to facilitate the morphological polarisation of newborn neurons through the initiation of a clearly defined axon. Notably, reducing the relative levels of ciliary GPR161 decreases ciliary cAMP levels and leads to disorganised actin aggregations, resulting in multiple, unstable axon-like projections of differentiating neurons. These findings underscore the importance of localised cAMP signalling in regulating neuronal polarity. Consistent with our observations, elevated ciliary cAMP levels, but not centrosomal cAMP, have been shown to modulate cell polarity during the migration of cortical interneurons^29^, highlighting the necessity of spatial and temporal regulation of this second messenger. Furthermore, vertebrate cells have been shown to distinguish between ciliary and extraciliary cAMP pools through the use of compartmentalised PKA, emphasising the importance of cAMP microdomains in the correct interpretation of signalling^30^.

Our results indicate that optimal ciliary cAMP levels, maintained by balanced ciliary GPR161 and Smo accumulation, define a transient cell state during which differentiating neurons become competent to polarise and initiate a nascent axon. These findings highlight the importance of ciliary cAMP in regulating the spatial organisation of actin, which leads to maturation of a singular actin accumulation into an axon, providing unexpected new insights into the regulation of actin dynamics through primary cilium-mediated non-canonical Shh signalling. However, the precise mechanisms through which ciliary cAMP levels modulate actin dynamics and its intermediate regulators remain unclear. Furthermore, given that current models of non-canonical Shh signalling are derived mainly from contexts in which signalling occurs outside the primary cilium^31,32^, a cilium-independent mode of non-canonical Shh transduction participating during neuronal differentiation cannot be excluded. Future studies exploring the role of ciliary and extraciliary Smo and actin-modulating proteins in this process will delineate the intermediate signalling steps linking primary cilium and non-canonical Shh signal transduction to the actin cytoskeleton. Although the crosslink between primary cilia and actin dynamics is not fully understood, it is well established that ciliary dysfunction disrupts actin organisation, leading to defects in cellular processes such as migration^33–35^. Moreover, increasing evidence suggests a reciprocal relationship between actin depolymerisation and ciliogenesis^36^, highlighting the potential for bidirectional regulation between the actin cytoskeleton and primary cilia.

Considering that GPR161 is normally excluded from the primary cilia upon exposure to Shh and activation of Smo^7^, it is important to investigate the mechanisms that facilitate the atypical concurrent GPR161 and Smo ciliary accumulation in newborn neurons. On a broader scale, given the nature of cAMP as a second messenger with diverse regulatory roles, it is likely that ciliary cAMP in the remodelled primary cilium leads to a broad range of downstream effects. As the cAMP signalling pathway has been reported to play a cooperative role with Ca^2+^ signalling in eukaryotic cells ^37,38^, and Ca^2+^ signalling at the primary cilium regulates Shh signalling^39,40^, it would be of particular interest to determine the crosstalk between cAMP and Ca^2+^ signalling in the remodelled primary cilium.

Finally, this work may also shed light upon the mechanistic basis for the manifestation of ciliopathies such as Joubert and Meckel syndromes, which are characterised by brain malformations caused by errors in axon navigation^41^. These have recently been hypothesised to be caused by inappropriate interpretation of Shh signalling^42^. Indeed, Joubert syndrome patient-derived cells display increased entry of Smo and more rapid exit of GPR161 from the cilium, resulting in an imbalance of these signalling components that leads to inappropriate activation of canonical Shh signalling^43^. The findings presented here provide a potential mechanistic explanation to this model, as it is now evident that differentiating neurons adjust their ciliary composition to reinterpret Shh signalling to achieve the correct sequence of cell state transitions that ultimately facilitate the formation of functional neural architecture. Since ciliopathies are multi-syndromic and often impact multiple organ systems, it is now important to investigate whether development of other tissue types is driven by similar primary cilium remodelling events.

## Materials and Methods

### In ovo electroporation and plasmids

Fertilised chicken (Gallus gallus domesticus) eggs were obtained from Medeggs Ltd (Heath Top Farm Norwich Road Fakenham NR21 8LZ UK) and incubated at 37°C to Hamburger and Hamilton stage 10-12. Spinal cords were electroporated as described previously ^17^, using the minimal concentration of DNA that allowed live visualisation. Neurogenin-2 (25ng/ul) was added to promote neuronal differentiation. Plasmids were used at concentrations between 25ng/ul and 400ng/ul. Details of plasmids are shown in the following table:

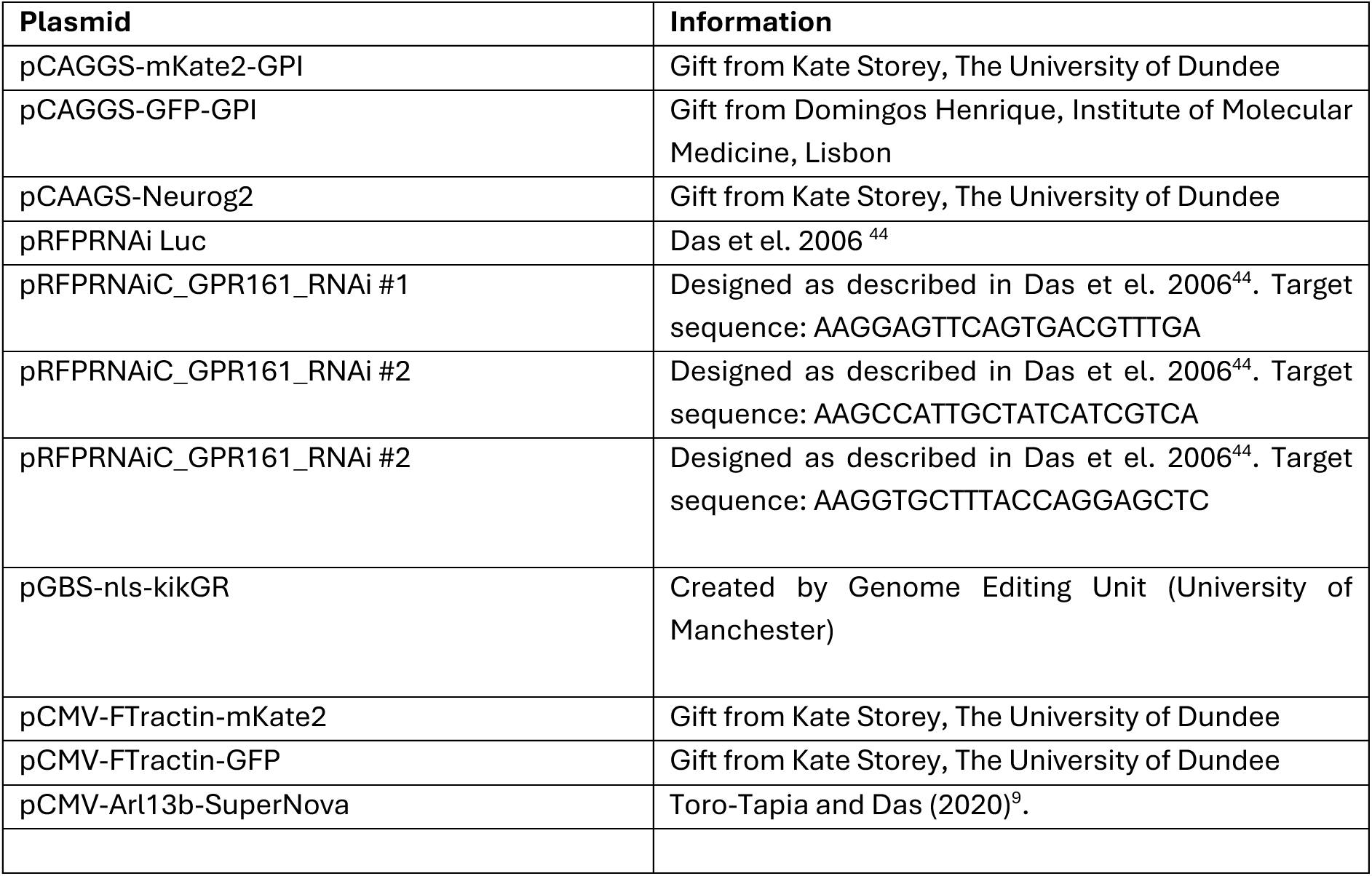

### Immunofluorescence, HCR RNA-FISH and fixed tissue imaging

Mouse embryos were fixed in 4% formaldehyde for 4 hours at 4 °C followed by dehydration in 30% sucrose in PBS overnight. These were then mounted in 1.5% LB agar / 30% sucrose and snap-frozen on dry ice. 20µm sections were collected using a Leica CM1950 cryostat and then processed for immunofluorescence. For immunofluorescence, samples were permeabilised with 0.1% Triton X-100 (Sigma-Aldrich) and blocked in 1% donkey serum (Sigma). Immunostaining requiring the use of primary antibodies from the same host species was performed using the Flexable Coralite-plus antibody labelling kit (Coralite plus 488, Cat No. KFA001; Coralite Plus 555, Cat No. KFA002; and Coralite Plus 647, Cat No. KFA003; Proteintech), using double the amount of standard reaction volume for each antibody as recommended by the manufacturer to increase signal/background fluorescence. Primary antibodies were used at the following dilutions: Tuj1 (80120, Biolegend) 1:1000, Arl13b (17711-1-AP, Proteintech) 1:100, Smo (20787-1-AP, Proteintech) 1:100 and GPR161 (13398-1-AP, Proteintech) 1:100. Secondary mouse antibody 405 Alexa Fluor (A48257 Invitrogen) was used at 1:250. Samples were mounted using ProLong™ Diamond Antifade Mountant (P36970, Thermo Fisher Scientific). For HCR RNA-FISH (Molecular Instruments), the probes for GRP161 were custom-designed using the first 1317 bp (from the start codon) shared among the nine predicted transcript sequences from Gallus gallus shown in NCBI (Accession: XM_040660182.2, XM_040660181.2, XM_040660183.2, XM_040660185.2, XM_040660187.2, XM_040660186.2, XM_040660188.2, XM_046905474.1, XM_046905493.1). Frozen tissue sections were processed for HCR according to the manufacturer’s protocol. Images were acquired using a Zeiss Cell Observer Z1 microscope system (Carl Zeiss) equipped with a 60x 1.40 NA objective, a Colibri 7 LED light source and a Flash4 v2 sCMOS camera (Hamamatsu).

### Embryo slice culture and time-lapse imaging

Embryos were electroporated as described earlier, incubated overnight, sliced and embedded in collagen I (Corning, Cat No. 354236) on a glass-bottomed dish (FluoroDish, World Precision Instruments, Cat No. FD35PDL-100) as described in detail previously^17^. Slices were taken from the trunk region between the wing and leg buds. Embedded slices were allowed to recover for an hour at 37°C in a 5% CO2 incubator in Neurobasal medium (ThermoFisher, Cat No 2348017) supplemented with B27 (ThermoFisher Scientific, 17504-044) and Glutamax (Gibco) before timelapse imaging. For inhibition experiments, the medium was replaced with medium containing cyclopamine (15 μM, Stratech Scientific) or ethanol (Merck), SAG (20 μM Cayman Chemical), Forskolin (Frk 10 μM, Stratech Scientific), or DMSO (Molecular Probes™ D12345) at the start of imaging.

Slices were imaged using a Zeiss Cell Observer Z1 system enclosed in an environment chamber maintained at 37°C and 5% CO_2_ equipped with a 40x 1.2 NA silicone immersion objective (Carl Zeiss), a Colibri 7 LED light source (Carl Zeiss), and a Flash4 v2 sCMOS camera (Hamamatsu). Image stacks were captured with minimal exposure times ranging from 20 to 50 milliseconds for each channel. Image stacks comprised of 40 to 50 optical sections, spaced 1.5 µm apart, with intervals of 5 minutes (for F-actin labelling) or 10 minutes between exposures. Imaging was conducted over a duration of 5 to 24 hours. Live imaging of actin dynamics was also performed using the Elyra 7 lattice structured illumination microscopy (SIM) system (Zeiss). Images were acquired every 2 minutes over a 4-hour imaging period using a 40x 1.2 NA water autocorrection FCS objective (Zeiss) and the AxioCam 820 camera (Zeiss), with a laser exposure time of 10 ms and low laser intensity (488 nm at 5%; 561 nm at 10%), and 2x binning. This was processed using the SIM2 algorithm with automatic sharpness and normalisation.

### Image processing

Deconvolution and post-processing were performed in Zen 2.3 (Blue edition, Carl Zeiss). Images acquired at 1×1 binning were deconvolved using the constrained iterative algorithm for a maximum of 40 iterations or until a 0.1% quality threshold was achieved. Images acquired with a 2×2 binning were deconvolved using the fast-iterative algorithm (Richardson-Lucy likelihood), applying the same number of iterations and quality threshold.

### Chromophore-assisted light inactivation

For chromophore-assisted light inactivation (CALI) experiments, E2 embryos were electroporated with F-Tractin-GFP, Neurog2 and Arl13b-SuperNova and ex vivo slice culture was performed as described previously. Sustained green light irradiation was carried out by capturing a Z-stack of 40 sections, each spaced 1.5 μm, over 10 seconds at 100 % intensity. Each cell was subjected to three cycles of green light irradiation, totalling 20 minutes of exposure. To confirm the loss of red fluorescence, an additional single Z-stack image of F-Tractin-GFP and Arl13b-SuperNova was taken for each cell of interest. Timelapse imaging was then performed to image F-Tractin-GFP every 5 minutes over a period of up to 16 hours. Control experiments performed included: imaging F-Tractin-GFP every 5 minutes over a period of up to 16 hours in cells transfected with F-Tractin-GFP, Arl13b-SuperNova and Neurog2 not subjected to sustained green light irradiation, and imaging F-Tractin-GFP every 5 minutes over a period of up to 16 hours in cells subjected to sustained green light irradiation but transfected with F-Tractin and Neurog2 only.

### Cilium-targeted cADDis and measurements

Ratiometric Cilium-Targeted cADDis mRNA codon optimised for chicken was obtained from Montana Molecular, Bozeman, Montana 59715, USA) and transfected into cells by *in ovo*-electroporation. Measurement of ciliary cADDis fluorescence intensity was performed in Zen Blue by tracing the contour of each primary cilium over time in a single plane. Mean fluorescence intensity was recorded and normalised by subtracting the background. Relative values were obtained by normalising with the mean fluorescence intensity from the first time point observed and graphed using GraphPad Prism version 10.0 (GraphPad Software, Boston, Massachusetts, USA). The Area Under the Curve (AUC) was calculated, and comparisons were performed using an unpaired t-test (GPR161 RNAi and SAG experiments) or a Mann-Whitney test (Forskolin experiment) in GraphPad Prism.

## Funding

Medical Research Council project grant MR/X008363/1 (RMD).

Biotechnology and Biological Sciences Research Council studentship 2441389 (RMD, HBB).

## Author contributions

Conceptualisation: RMD and GT-T.

Methodology: GT-T, HBB and RMD.

Investigation: GT-T and HBB.

Formal analysis: GT-T, HBB, JRD and VB.

Supervision: RMD

Writing – original draft: RMD and GT-T

Writing – review and editing: RMD, GT-T, HBB, VB, JRD.

## Data and materials availability

All data needed to evaluate the conclusions in the paper are present in the paper and/or the Supplementary Materials. Raw data files and all reagents may be requested from the authors. Cilium-targeted cADDis was used under MTA from Montana Molecular.

