## Supplementary Information for "Ciliary cAMP regulates Shh signal interpretation to drive polarisation of differentiating neurons"

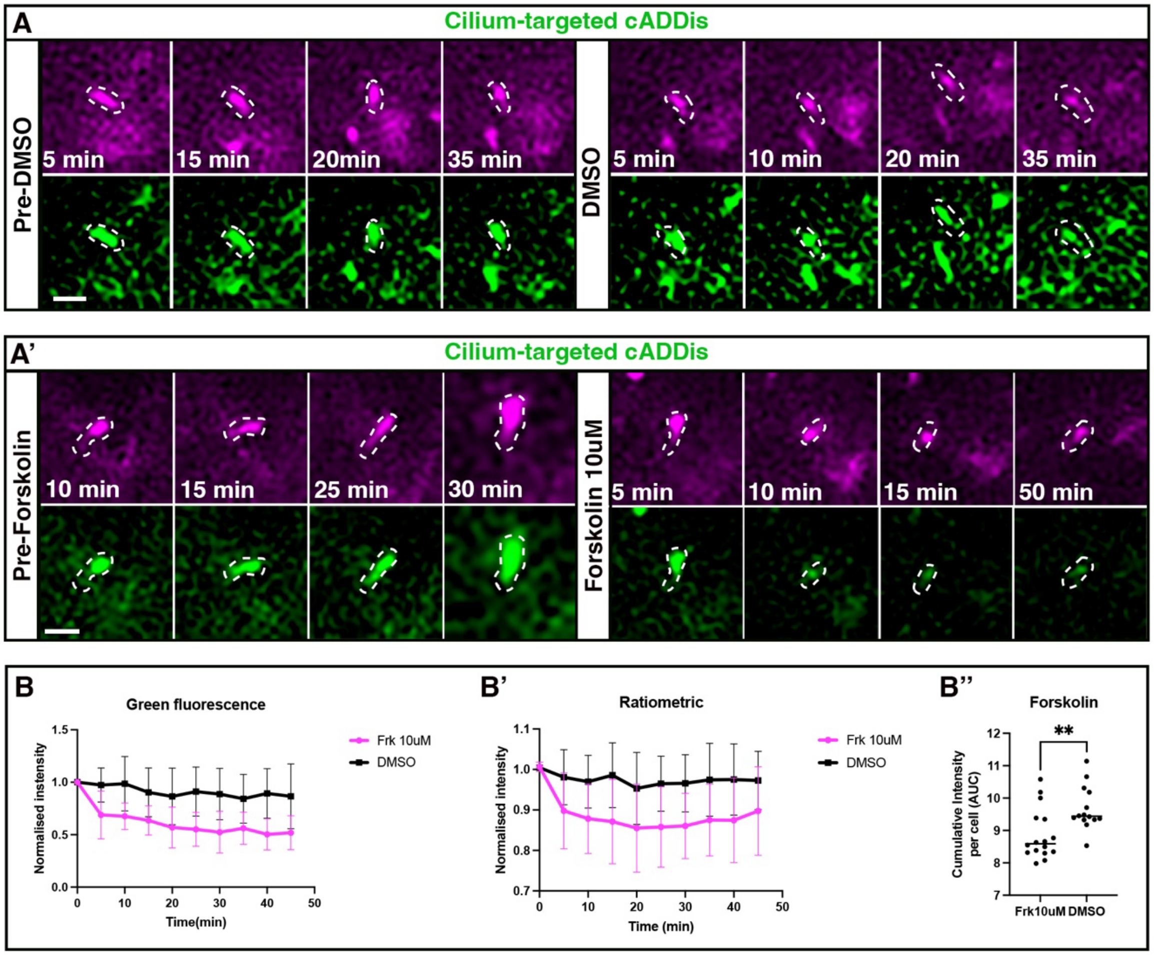


**Figure S1. Validation of cilium-targeted cADDis reporter activity. (A)** Timelapse sequence of neuroepithelial progenitor cell primary cilium before and after application of DMSO to imaging medium. The cell expresses a ratiometric, cilium-targeted 5-HT6-mCherry-cADDis sensor (green) with an mCherry fluorescent tag for ratiometric measurement (magenta). **(A')** Timelapse sequence of neuroepithelial progenitor cell primary cilium before and after application of 10 µM forskolin (Fsk) to imaging medium. **(B)** Quantification of normalised green ciliary fluorescence of cells imaged in medium containing DMSO (black line) or Fsk (magenta). Error bars represent standard deviation. **(B')** Measurement of ratiometrically normalised green ciliary fluorescence of cells imaged in medium containing DMSO (black line) or Fsk (magenta). Error bars represent standard deviation. **(B'')** Quantification of cumulative intensity per cell (AUC, area under the curve). Data analysed using a non-parametric Mann-Whitney test ($p=0.046$). Scale bars: 2µm.

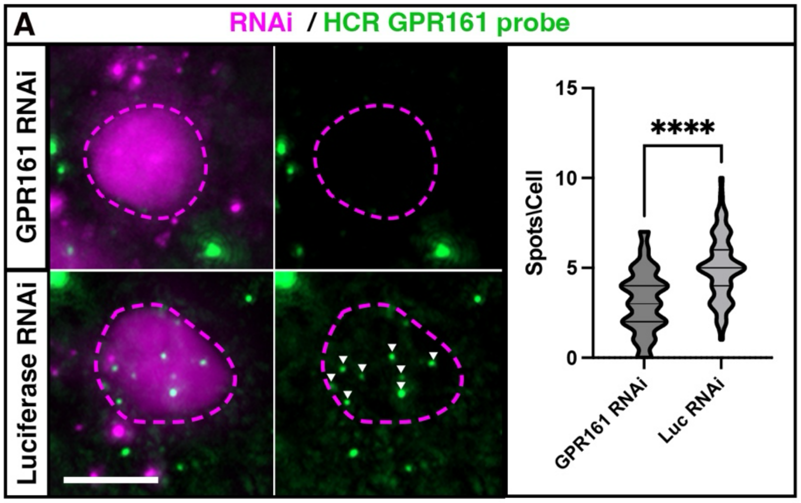


**Figure S2. Validation of GPR161 knockdown. (A)** HCR RNA-FISH assay to detect GPR161 mRNA expression (green dots) in cells transfected with GPR161 RNAi constructs (top panels) or a Luciferase RNAi construct (bottom panels). Magenta dashed lines indicate cell bodies. Graph displays quantification of knockdown efficiency. Data analysed using unpaired t-test ($p < 0.0001$). Scale bars: 5µm.

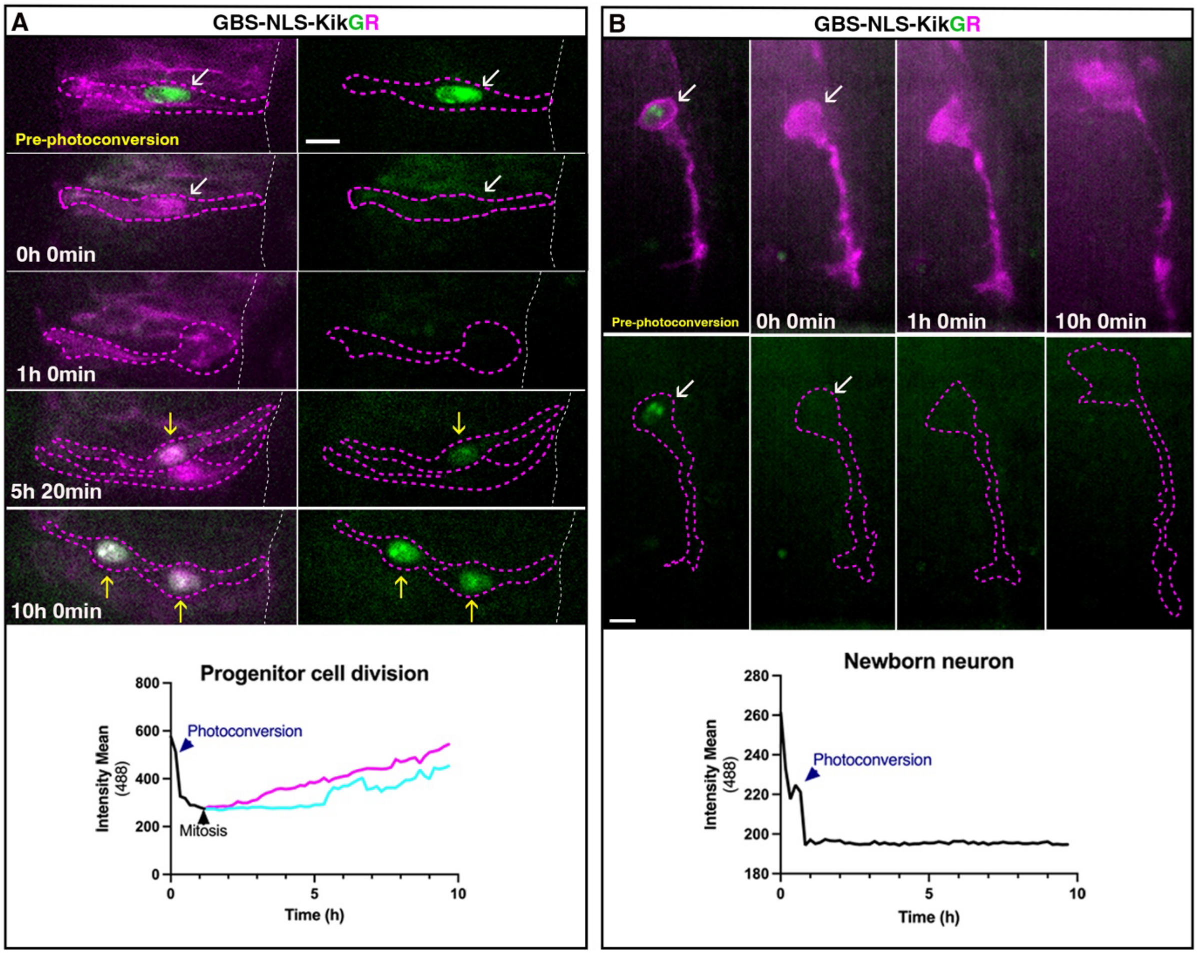


**Figure S3. Validation of photoconvertible reporter for Gli transcriptional activity GBS-NLS-KikGR. (A)** Timelapse sequence of neural progenitor cell expressing the green-to-red photoconvertible Gli reporter GBS-NLS-KikGR and mKate2-GPI (magenta) undergoing cell division. Top panels show cell before photoconversion. Magenta dashed lines demarcate cell membrane. White arrows indicate expression of nuclear KiKGR before and after photoconversion. Yellow arrows indicate recovery of green fluorescence in nuclei of daughter cells. The bottom graph displays quantification of mean fluorescence intensity for the green channel from this representative timelapse sequence. **(B)** Timelapse sequence of differentiating neuron expressing the green-to-red photoconvertible Gli reporter GBS-NLS-KikGR and mKate2-GPI (magenta) undergoing axon extension. Left-hand panels show cell before photoconversion. White arrows indicate nucleus and magenta dashed lines demarcate the cell membrane. Bottom graph displays quantification of mean fluorescence intensity for the green channel (488) over time from this representative timelapse. Scale bars: 10µm.

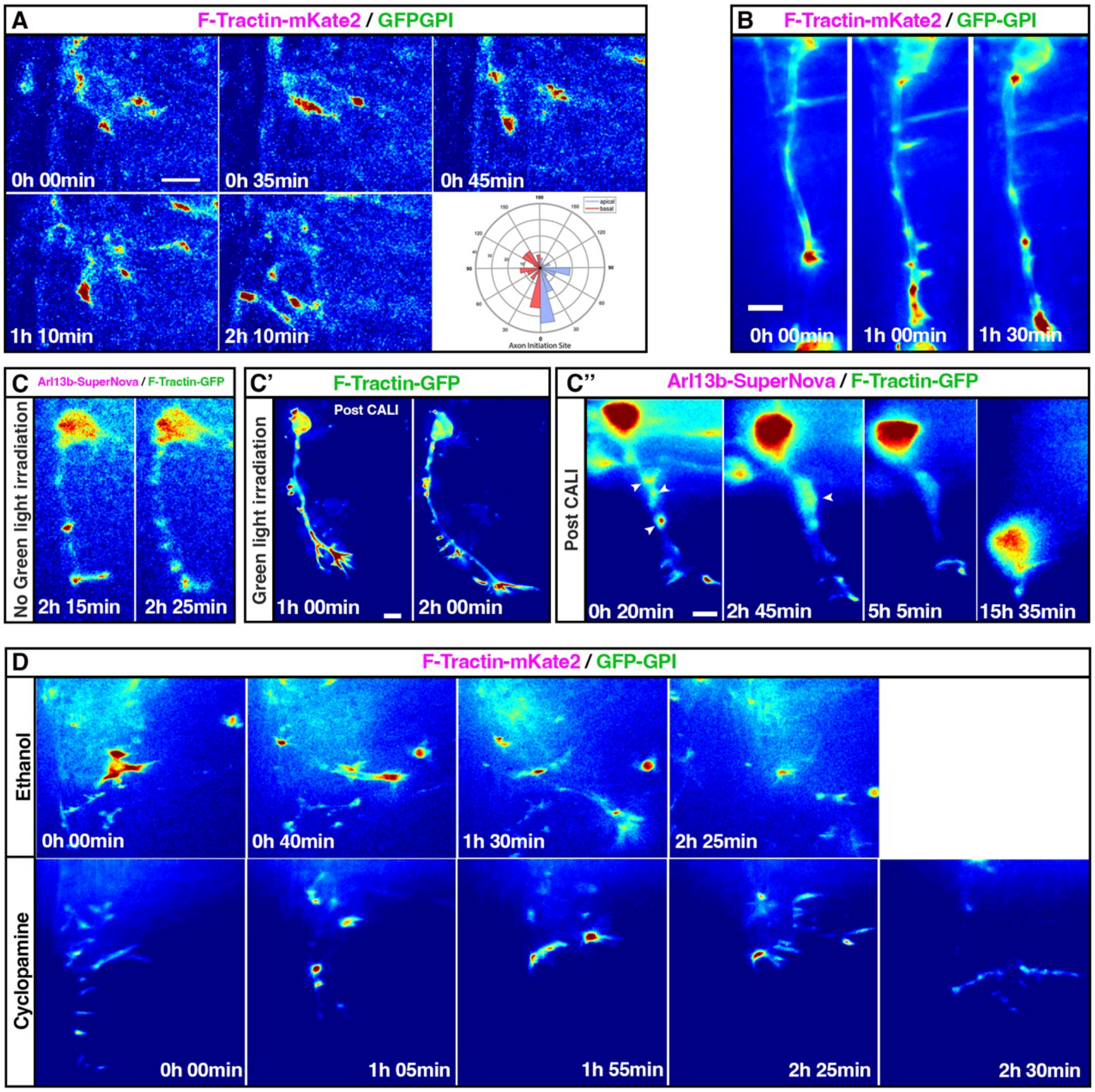


**Figure S4. Primary cilium-mediated Shh signal interpretation facilitates normal actin dynamics in differentiating neurons.** This figure mirrors experiments from Figure 4 with the F-Tractin channel displayed using the jet colour map to highlight actin accumulations. **(A)** Timelapse sequence of differentiating neuron expressing F-Tractin-mKate2 and GFP-GPI undergoing axon initiation. Rose plot of the positions of F-actin accumulations that mature into initiating axon across 25 cells with respect to the axon initiation site. **(B)** Timelapse sequence of cell expressing F-Tractin-mKate2 and GFP-GPI undergoing axon extension. **(C)** Timelapse sequence of cell expressing Arl13b-SuperNova and F-Tractin-GFP undergoing normal axon extension while not subjected to green-light irradiation. **(C')** Timelapse sequence of cell expressing F-Tractin-GFP only undergoing normal axon extension following green-light irradiation. **(C'')** Timelapse sequence of cell expressing Arl13b-SuperNova and F-Tractin-GFP subjected to green-light irradiation. **(D)** Timelapse sequence of cell expressing F-Tractin-mKate2 and GFP-GPI undergoing axon extension imaged in medium containing vehicle control ethanol (top panels) and cyclopamine (bottom panels). Scale bars: 10μ m.

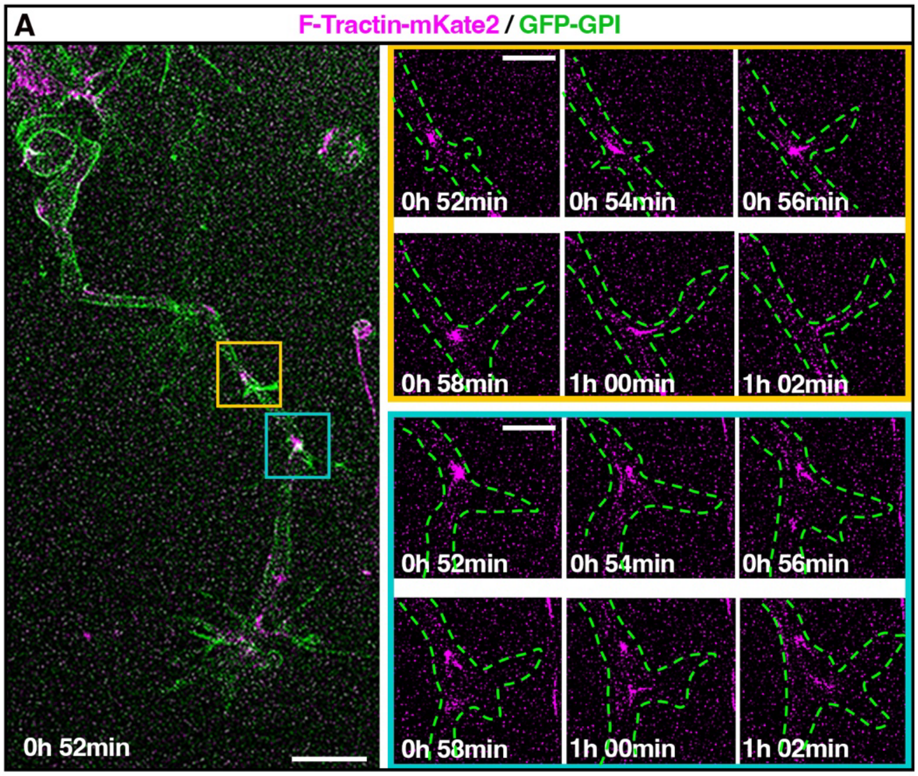


**Figure S5. Axon extension is characterised by dynamic actin-rich axonal protrusions. (A)** Structured illumination microscopy timelapse sequence of axonal actin accumulations (yellow and cyan boxes) in cell expressing F-Tractin-mKate2 (magenta) and GFP-GPI (green) undergoing axon extension. Enlarged panels highlight axonal F-actin accumulations that extend into the axonal protrusion. Scale bars, 10µm (left panel) and 5µm (zoomed-in right panels).

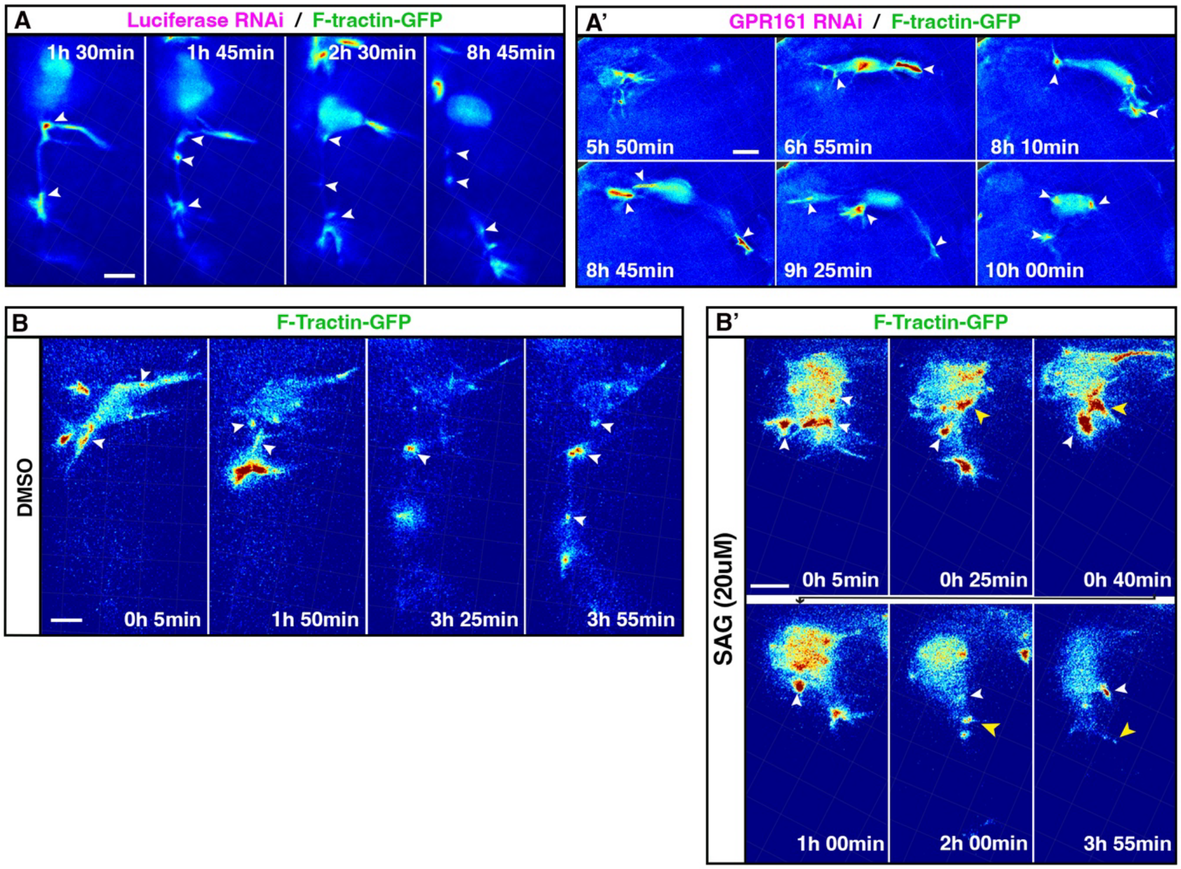


**Figure S6. Ciliary cAMP levels direct actin dynamics in differentiating neurons.** This figure mirrors experiments from Figure 5 with the F-Tractin channel displayed using the jet colour map to highlight actin accumulations. **(A)** Timelapse sequence of differentiating neuron expressing luciferase RNAi construct and F-Tractin-GFP. White arrowheads indicate dynamic axonal actin accumulations. **(A')** Timelapse sequence of differentiating neuron expressing GPR161 RNAi constructs and F-Tractin-GFP. White arrowheads indicate actin aggregation in unstable axon-like projections. **(B)** Timelapse sequence of a differentiating neuron expressing F-Tractin-GFP imaged in medium containing DMSOWhite arrowheads indicate dynamic axonal actin accumulations. **(B')** Timelapse sequence of differentiating neuron expressing F-Tractin-GFP imaged in medium containing 20 µM SAG. White arrowheads indicate actin aggregations in cell body and yellow arrowheads indicate unstable axon-like projections. Scale bars: 10µm.

### Movie Legends

**Movie S1: Relative cAMP levels increase in the remodelled primary cilium of differentiating neurons**

Timelapse sequence of a differentiating neuron undergoing apical process retraction expressing cilium-targeted cADDis (green) and membrane marker mKate2-GPI (magenta).

**Movie S2: Expression of a luciferase-targeting shRNA construct does not affect axon extension.**

Timelapse sequence of differentiating neuron expressing luciferase-targeting RNAi construct (magenta) and membrane marker GFP-GPI (green).

**Movie S3: GPR161 knockdown disrupts axon initiation.**

Timelapse sequence of differentiating neuron expressing GPR161-targeting RNAi constructs (magenta) and GFP-GPI (green).

**Movie S4: Elevated relative ciliary cAMP levels in differentiating neuron expressing luciferase-targeting shRNA construct.**

Timelapse sequence of differentiating neuron expressing luciferase RNAi construct (magenta) and cilium-targeted cADDis (green).

**Movie S5: Decreased relative ciliary cAMP levels in differentiating neuron expressing GPR161 knockdown construct.**

Timelapse sequence of differentiating neurons expressing GPR161 RNAi constructs (magenta) and cilium-targeted cADDis (green).

**Movie S6: Elevated relative ciliary cAMP levels in differentiating neuron imaged in medium containing DMSO.**

Timelapse sequence of differentiating neuron expressing cilium-targeted cADDis (green) and mKate2-GPI (magenta) imaged in medium containing DMSO

**Movie S7: Decreased relative ciliary cAMP levels in differentiating neuron cultured in medium containing SAG.**

Timelapse sequence of differentiating neuron expressing cilium-targeted cADDis (green) and mKate2-GPI (magenta) imaged in medium containing 20 μM SAG

**Movie S8: Validation of GBS-KIKGR in neural progenitors undergoing cell division.**

Timelapse sequence of neural progenitor cell expressing the green-to-red photoconvertible Gli reporter GBS-NLS-KikGR and mKate2-GPI (magenta) undergoing cell division

**Movie S9: Validation of GBS-KIKGR in differentiating neurons undergoing axon extension.**

Timelapse sequence of differentiating neuron expressing the green-to-red photoconvertible Gli reporter GBS-NLS-KikGR and mKate2-GPI (magenta) undergoing axon extension

**Movie S10: Differentiating neuron expressing luciferase-targeting RNAi switches to non-canonical Shh signalling.**
Timelapse sequence of differentiating neuron expressing luciferase RNAi construct (magenta) and GBS-NLS-KikGR

**Movie S11: Maintenance of canonical Shh signalling in differentiating neuron expressing GPR161 knockdown constructs**.
Timelapse sequence of differentiating neuron expressing GPR161 RNAi constructs (magenta) and GBS-NLS-KikGR.

**Movie S12: Differentiating neuron imaged in medium containing DMSO undergoes the switch to non-canonical Shh signalling.**
Timelapse sequence of differentiating neuron expressing GBS-NLS-KikGR and membrane marker mKate2-GPI (magenta) imaged in medium containing DMSO.

**Movie S13: Activation of Gli transcriptional activity in differentiating neuron imaged in medium containing SAG.**
Timelapse sequence of differentiating neuron expressing GBS-NLS-KikGR and mKate2-GPI (magenta) imaged in medium containing 20 μM SAG.

**Movie S14: Axon initiation is preceded by localised F-actin accumulation.**
Timelapse sequence of differentiating neurons expressing F-actin probe F-Tractin-mKate2 (magenta) and membrane marker GFP-GPI (green) undergoing axon initiation.

**Movie S15: Axon extension is characterised by dynamic actin-rich axonal protrusions.**
Timelapse sequence of differentiating neuron expressing F-Tractin-mKate2 (magenta), and GFP-GPI (green) undergoing axon extension

**Movie S16: F-actin accumulations are present within the axon shaft and appear to migrate into the axonal protrusion**
Structured illumination microscopy timelapse of axonal actin accumulations in cell expressing F-Tractin-mKate2 (magenta) and GFP-GPI (green) undergoing axon extension.

**Movie S17: Normal actin dynamics in CALI control not subjected to green-light irradiation.**
Timelapse sequence of differentiating neuron expressing Arl13b-SuperNova (magenta) and F-Tractin-GFP (green), not subjected to green-light irradiation, undergoing normal axon extension.

**Movie S18**: **Normal actin dynamics in CALI control subjected to green-light irradiation.**
Timelapse sequence of differentiating neuron expressing F-Tractin-GFP (green), subjected to green-light irradiation, undergoing normal axon extension.

**Movie S19: Disruption of primary cilium function inhibits normal actin dynamics and leads to axon collapse.**
Timelapse sequence of differentiating neuron expressing Arl13b-SuperNova (magenta) and F-Tractin-GFP (green) following CALI-mediated disruption of primary cilium.

**Movie S20: Carrier ethanol does not alter actin dynamics.**
Timelapse sequences of differentiating neurons expressing F-Tractin-mKate2 (magenta), and GFP-GPI (green) undergoing axon extension, imaged in medium containing ethanol.

**Movie S21: Cyclopamine mediated inhibition of Smo function alters actin dynamics and increases filopodia-like protrusions.**
Timelapse sequences of differentiating neurons expressing F-Tractin-mKate2 (magenta), and GFP-GPI (green) undergoing axon extension imaged in medium containing 15 μM cyclopamine

**Movie S22: Differentiating neuron expressing luciferase RNAi construct presents normal actin dynamics**
Timelapse sequence of differentiating neuron expressing luciferase RNAi construct (magenta) and F-Tractin-GFP (green) to label filamentous actin.

**Movie S23: Differentiating neuron expressing GPR161 RNAi constructs presents unstable axon-like projections that emerge from aggregated actin accumulations within the cell body.**
Timelapse sequence of differentiating neuron expressing GPR161 RNAi constructs (magenta) and F-Tractin-GFP (green).

**Movie S24: Normal actin dynamics in differentiating neuron imaged in medium containing DMSO.**
Timelapse sequence of differentiating neuron expressing F-Tractin-GFP (green) imaged in medium containing DMSO.

**Movie S25: Differentiating neuron imaged in a medium containing SAG presents unstable axon-like projections that emerge from aggregated actin accumulations within the cell body.**
Timelapse sequence of differentiating neuron expressing F-Tractin-GFP (green) imaged in medium containing 20 μM SAG.
